# Trial-by-trial fluctuations in amygdala activity track motivational enhancement of desirable sensory evidence during perceptual decision-making

**DOI:** 10.1101/2021.12.03.471135

**Authors:** Ren Paterson, Yizhou Lyu, Yuan Chang Leong

## Abstract

People are biased towards seeing outcomes that they are motivated to see. For example, sports fans often perceive the same ambiguous foul in favor of the team they support. Here, we test the hypothesis that motivational biases in perceptual decision-making arise from amygdala-dependent biases in sensory processing. Human participants were rewarded for correctly categorizing an ambiguous image into one of two categories while undergoing fMRI. On each trial, we used a financial bonus to motivate participants to see one category over another. The reward maximizing strategy was to perform the categorizations accurately, but participants were biased towards categorizing the images as the category we motivated them to see. Heightened amygdala activity was associated with motivation consistent categorizations, and tracked trial-by-trial enhancement of neural activity in sensory cortices that was specific to the desirable category. Analyses using a drift diffusion model provide converging evidence that trial-by-trial amygdala activity was associated with stronger biases in the accumulation of sensory evidence. Prior work examining biases in perceptual decision-making have focused on the role of frontoparietal regions. Our work highlights an important contribution of the amygdala. When people are motivated to see one outcome over another, the amygdala biases perceptual decisions towards those outcomes.

---

People are biased towards seeing percepts that they are motivated to see, a phenomenon known as ‘wishful seeing’ (Dunning and Balcetis 2013). For example, sports fans of opposing teams watching the same game often perceive an ambiguous foul differently, with each group of fans judging the foul in favor of the team they support. When motivated to see one interpretation of reality over another, people become less objective and are more prone to perceptual errors (Voss et al. 2008; Bromberg-Martin and Sharot 2020; Leong et al. 2021). Individuals who score higher on trait paranoia are also more likely to exhibit wishful seeing (Rossi-Goldthorpe et al. 2021), suggesting that extreme cases of wishful seeing might contribute to biased beliefs about the world that are the hallmark of many psychiatric disorders (Kube and Rozenkrantz 2021). Why are people more likely to see what they are motivated to see?

To formulate a mechanistic understanding of wishful seeing, we can draw on the extensive neurophysiological and behavioral modeling work characterizing how perceptual decisions are determined from noisy sensory information (e.g., Heekeren et al., 2008; Shadlen and Kiani, 2013). In particular, perceptual decisions are thought to involve the temporal accumulation of sensory evidence from the environment. When the level of sensory evidence reaches a threshold, the individual commits to a decision. Recent work suggests that wishful seeing is driven by the selective enhancement of neural activity encoding desirable percepts (Leong et al. 2019). This biases sensory evidence accumulation in favor of percepts one is motivated to see, giving rise to motivationally biased perceptual decisions. While prior work has shown that motivation enhances the neural representation of desirable percepts in sensory cortices, how this enhancement occurs is not known. Consequently, the neural mechanisms that give rise to wishful seeing remain underspecified.

The amygdala has extensive connections to the visual cortex (Freese and Amaral 2005), and is well-positioned to mediate wishful seeing. Human neuroimaging and non-human primate electrophysiology studies have found that the amygdala plays a central role in biasing attention towards affectively salient aspects of the environment (Pessoa and Adolphs 2010; Pourtois et al. 2013; Peck and Salzman 2014a; Mather et al. 2016). For example, fluctuations in amygdala activity correlate with the enhanced neural representation of images that had acquired affective salience due to having been previously paired with an electrical shock (Lim et al. 2009). We hypothesize that wanting to see a percept ascribes affective significance to the associated sensory features, and the amygdala is similarly involved in enhancing the neural representation of these features. Past studies, however, have not examined the role of the amygdala in contexts where participants are motivationally biased to see one percept over another. It is thus unclear if and how the amygdala biases perception towards desirable percepts.

In the current work, we conducted a series of pre-registered analyses on a previously published dataset (Leong et al. 2019) to test the hypothesis that the amygdala facilitates wishful seeing by enhancing sensory evidence accumulation in favor of desirable percepts. We first assessed if amygdala activity was associated with motivationally biased perceptual decisions. Next, we tested if amygdala activity was associated with stronger neural representations of percepts participants were motivated to see. Finally, we incorporated participants’ trial-by-trial amygdala activity, perceptual decisions and response times into a computational model to test the hypothesis that amygdala activity was specifically associated with motivational biases in evidence accumulation. Together, our work takes a convergent approach that combines behavioral, neural and modeling measures to characterize the neural mechanisms underlying motivational biases in perceptual decision-making.

## Materials and Methods

### Participants

Thirty-three participants provided informed consent and were compensated between $30-50 depending on their performance in the task. Data were discarded from 3 participants due to excessive head motion (i.e., >3 mm in any direction) during >1 scanning sessions, yielding a final sample of 30 participants (17 males, 13 females; age range, 18-43 years; mean age, 22.3 years). These data have been previously reported in Leong et al. (2019).

### Stimuli

Stimuli consisted of grayscale images created by combining a scene image and face image in varying proportions. Face images were frontal photographs taken from the Chicago Face Database (Ma et al. 2015), and consisted of half female and half male faces with neutral expressions. Scene images consisted of half outdoor and half indoor scenes. For each participant, six stimulus sets were created. Each set contained 40 images: 1 × 100% scene, 3 × 65% scene, 5 × 60% scene, 7 × 55% scene, 8 × 50% scene, 7 × 45% scene, 5 × 40% scene, 3 × 35% scene and 1 × 0% scene. Images with more than 50% scene were considered “more scene” while images with less than 50% scene were considered “more face”. Stimuli were presented to participants using MATLAB (Mathworks) and the Psychophysics Toolbox (Brainard 1997).

### Experimental Task

Participants were told that they would be performing a visual categorization task with a teammate (“Cooperation” condition) and an opponent (“Competition” condition) (Fig. 1). At the beginning of each trial, either the teammate or the opponent would make a bet on whether the upcoming image would consist of more face or more scene (i.e. whether the face image or scene image was of higher intensity). Participants were first presented with the text “Your Teammate/Opponent bets 40 cents that the next image will have more” for 1 second (*Trial Start)*, before the category that the teammate/opponent bet on was shown on screen for 3 seconds (*Motivation Cue*). Participants were then presented with a face-scene composite image (see *Stimuli*). Participants earn 40 cents if the teammate’s bet was correct (e.g., if the teammate bet “more face” and the image actually consists of more face) and lose 40 cents if the teammate’s bet was wrong. In contrast, participants lose 40 cents if the opponent’s bet was correct and earn 40 cents if the opponent’s bet was wrong. As such, participants were motivated to see the image as the category that the teammate had bet on and the category that the opponent had bet against.

**Figure 1.**
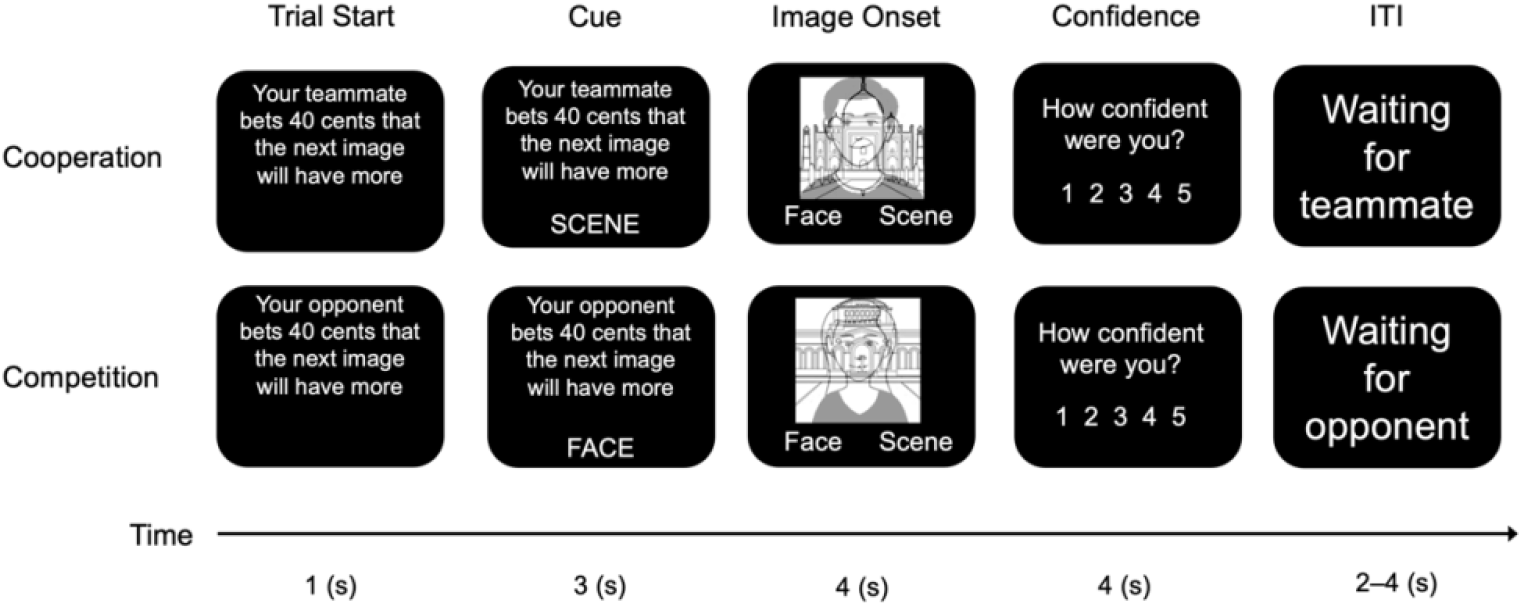
Experimental design. Each trial begins with a teammate (Cooperation condition) or an opponent (Competition condition) making a bet on whether the upcoming image would consist of more face or more scene. Participants earn 40 cents if the teammate’s bet is correct, and lose 40 cents otherwise. Conversely, they lose 40 cents if the opponent’s bet is correct and earn 40 cents otherwise. Participants are then presented with a face-scene composite image, and have to categorize whether the image has more face or more scene. They then rated how confident they were in their categorization. Images have been replaced with cartoon images to comply with *bioRxiv* policy.

Participants were then asked to indicate whether the image consists of more face or more scene and received a 10-cent reward if they correctly categorized the image. If participants did not respond in 4 seconds, the trial would time out and participants would lose the opportunity to earn the reward. The image remained on screen for the entire 4 seconds regardless of when participants made their response. Participants then indicated how confident they were in their categorization on a 1-5 scale. After a variable inter-trial-interval (ITI) of 2-4 seconds, they proceeded to the next trial. Participants performed 2 fMRI runs of the Cooperation condition and 2 fMRI runs of the Competition condition in an interleaved order that was counterbalanced across participants. Each run was approximately 8 min long and consisted of 40 trials.

Importantly, the 40-cent bonus associated with the teammate or opponent’s bet depended solely on the objective category of the image (i.e. whether the image actually contained more face or more scene) and was not affected by participants’ responses. For example, if the teammate bet that the upcoming image contains more face but the image objectively contained more scene, participants would lose the 40 cents regardless of how they categorized the image. As such, participants would earn the most amount of money from the experiment if they ignored the bets and categorized the images accurately based on what they saw to earn the maximum reward from correctly categorizing each image. Participants were told that both the teammate and the opponent did not have *a priori* information about the images and were making blind bets. Participants were also told that neither the teammate or opponent would be informed of their responses, and were thus not pressured to conform to the bets. Unbeknownst to participants, the teammate’s and opponent’s bets were pseudo-randomized to be correct on half of the trials.

The instructions were delivered verbally by the experimenter and also in written form on screen at the beginning of the experiment. It was intuitive to participants that the bet depended on the objective category of the images rather than their categorizations, and that they should categorize each image accurately to earn maximum reward. In the post-experiment survey, all participants responded that the task instructions were clear and that they had understood the task correctly.

### MRI Data Acquisition and Preprocessing

A 3T General Electric scanner was used to collect the MRI data. Functional images were obtained with T2*-weighted echo planar imaging (EPI) pulse sequences. Each volume comprised 46 transverse slices. Volumes were acquired in interleaved order, with the following imaging parameters: repetition time, 2 s; echo time, 25 ms; flip angle, 77°; voxel size, 2.9 *mm*^*3*^. A T1-weighted pulse sequence was used to acquire anatomical images at the beginning of the session with the following parameters: repetition time, 7.2 ms; echo time, 2.8 ms; flip angle, 12°; voxel size, 1*mm*^*3*^.

Preprocessing and analysis of EPI images were performed using FSL/FEAT v6.05 (FMRIB software library, FMRIB, Oxford, UK) and included motion correction (rigid-body realignment of functional volumes to the first volume), slice-timing correction, high-pass filtering of the data with a 100 ms cut-off, and spatial smoothing using a Gaussian kernel with a full-width at half-maximum of 4 mm. For multivoxel classification analyses, our classifier was trained and tested in each participant’s native space (see fMRI analyses). For all other analyses, we first registered the functional volumes to participants’ anatomical images (6 d.f.) and then registered the volumes to a template brain in the Montreal Neurological Institute (MNI) space (affine transformation with 12 d.f.).

### Regions of interest (ROI) definition

We defined a bilateral amygdala ROI as the voxels that are estimated to have greater than 0.5 probability as being part of the amygdala according to the Harvard-Oxford Subcortical Structural Atlas. To measure sensory representations, we defined an occipito-temporal ROI by combining the bilateral occipital and temporal lobe masks from the Harvard-Oxford Structural Atlas. This mask includes the visual cortex as well as the ventral temporal areas important for object recognition (Grill-Spector 2003). We defined additional ROIs from an atlas derived from applying independent component analysis to resting-state data (Shirer et al. 2012). Specifically, the anterior insula and dorsal anterior cingulate cortex (dACC) ROIs were obtained from the set of regions labeled the “anterior salience network”, while the dorsolateral prefrontal cortex (DLPFC) and inferior parietal lobule (IPL) ROIs were obtained from the set of regions labeled the “executive control network”.

### fMRI Analyses

In earlier work, we had found that motivational bias and response times did not differ between the Competition and Cooperation conditions - participants were just as likely and quickly to categorize the image as the category that the teammate had bet on as they were to categorize it as the category the opponent had bet against (Leong et al. 2019). Therefore, in the current study, we collapsed across the two conditions based on whether the desirable category was face (i.e. when the teammate bet on more face or when the opponent bet on more scene) or scene (i.e. when the teammate bet on more scene or when the opponent bet on more face).

We ran a general linear model (GLM) analysis contrasting neural activity on motivation consistent trials (i.e. trials where participants categorized the image as the desirable category) against that on motivation inconsistent trials (i.e. trials where participants categorized the image as the less desirable category). We focused specifically on activity at the time of the motivation cue (i.e. 3 seconds prior to the image onset). The two trial types were modeled as separate regressors. These were created by convolving a stimulus function, consisting of an impulse response at the time of the motivation cue, with a Gamma hemodynamic response function (HRF).

As is conventional for analyses using FSL, we estimated a separate model for each run before aggregating across the four runs of a participant in a fixed effects analysis to obtain a statistical parametric map. This approach allows for the estimation of between-run variability for each participant. For each model, response times (convolved with HRF) and extended motion parameters (i.e. standard motion parameters, their temporal derivatives, and the squares of both the standard motion parameters and the temporal derivatives; not convolved with HRF) were included as covariates of no interest. We then used FSL’s *featquery* function to extract the average *z*-statistic of the contrast in the amygdala ROI for each participant, and ran a one-sample t-test against zero to assess if amygdala activity was higher on motivation consistent trials than on motivation inconsistent trials.

At the end of the experimental task, participants performed two runs of a localizer task where they were presented with unambiguous face and scene images one at a time (5 blocks of 15 images of each category; each image presented for 2 seconds with a 2-second interval). Participants had to indicate whether each face was male or female, and whether each scene was indoors or outdoors. We trained an L1-regularized logistic regression classifier (C = 1) to classify whether the participant was seeing a face or scene based on the patterns of activity in the occipito-temporal ROI. We then applied the trained classifier to the single-trial activation patterns in the experimental task. Specifically, all trials in a run were simultaneously estimated in a single GLM, with each trial modeled using a separate regressor consisting of an impulse response at the time of stimulus onset convolved with a HRF (i.e. a “Least-Squares All” model; Mumford et al., 2014). The beta weights of each trial were then used as inputs for the trained classifier. For each trial, the classifier outputs the probability that the participant was viewing a scene vs. a face based on the beta weights in the occipito-temporal ROI, thus providing us with a measure of the relative strength of scene vs. face selective neural activity for that trial.

We estimated amygdala activity on each trial using a Least-Squares All GLM that modeled each trial as a separate regressor created by convolving a stimulus function, consisting of an impulse response at the time of each motivation cue, with a HRF. The resulting beta weights obtained for each regressor were then averaged within the amygdala ROI as the estimate of anticipatory amygdala activity for that trial. A linear mixed effects model (LME Model M1; Table S1) was used to test the relationship between trial-by-trial anticipatory amygdala activity and sensory representations:

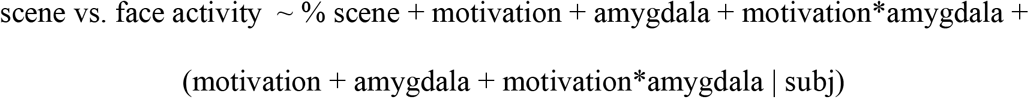

where *scene vs. face activity* is the relative strength of scene vs. face-selective neural activity as indexed by the classifier output, *% scene* is the amount of scene in an image, *motivation* is the category participants were motivated to see (face = 0, scene = 1), and *amygdala* is amygdala activation in response to the motivation cue. The coefficient for the interaction term, *motivation*amygdala*, would thus capture the extent to which motivational effects on sensory-related activity depended on anticipatory amygdala activation. To unpack the interaction effect, we divided each participant’s data into trials with high and low amygdala activity based on a median split, and examined the effect of motivation on *scene vs. face activity* for the two trial types separately (High Amygdala Activity: LME Model M2, Low Amygdala Activity: LME Model M3; Table S1):

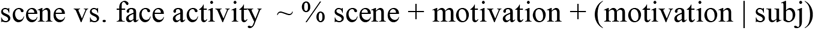

In a series of exploratory analyses, we also examined if motivational effects on scene and face-selective activity were dependent on activity in the anterior insula, dorsal anterior cingulate cortex (dACC), dorsolateral prefrontal cortex (DLPFC), and inferior parietal lobule (IPL) (Tables S2). All models were estimated using the lmer function in the lme4 package in R, with P values computed from t-tests with Satterthwaite approximation of degrees of freedom as implemented in the lmerTest package. Full model specification and estimates are reported in Table S1 and S2.

### Drift Diffusion Model

We fit a drift diffusion model to participants’ data using the HDDM toolbox with default priors (Wiecki et al. 2013). The model assumes that participants’ decisions are determined by the stochastic accumulation of sensory evidence towards one of two decision thresholds. We estimated model parameters that determined the starting point of the decision process (parameter *z*), the rate of evidence accumulation (i.e. *drift rate*, parameter *v*), the distance between the two decision thresholds (parameter *a*) and time not related to the decision process *(e*.*g*., time for stimulus encoding or motor response; *non-decision time*, parameter *t*). We modeled the “scene” decision threshold as the top boundary (i.e. scene threshold = *a*) and the “face” threshold as the bottom boundary (i.e. face threshold = *0*). HDDM implements hierarchical Bayesian estimation, which assumes that parameters for individual participants were randomly drawn from a group-level distribution. Individual participant parameters and group-level parameters were jointly estimated using Markov Chain Monte Carlo sampling (20,000 samples; burn-in=2,000 samples; thinning=2). To account for outliers generated by a process other than that assumed by the model (e.g., accidental button press, lapses in attention), we estimated a mixture model where 5% of trials were assumed to be distributed according to a uniform distribution. We chose not to include inter-trial variability parameters in this model, as simulation studies have shown that they can compromise the reliability and accuracy of the estimates of individual parameters (Boehm et al. 2018). A model that included parameters for inter-trial variability in the starting point, drift rate and non-decision time yielded qualitatively similar results (see *Supplemental Text*; Fig. S1).

The drift rate was allowed to vary trial-by-trial as a function of the stimulus, the category participants were motivated to see, and amygdala activity:

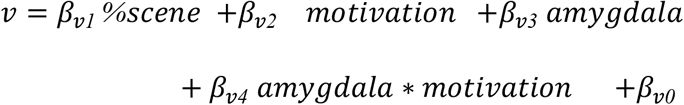

where *% scene* refers to the percentage scene in an image, *motivation* refers to the category participants were motivated to see (face = -1; scene = 1), and *amygdala* refers to amygdala activation at the onset of the motivation cue estimated from a GLM (see fMRI analysis). The coefficient for the *amygdala*motivation* interaction reflects the extent to which the effect of motivation on the drift rate depended on amygdala activation. Specifically, a positive value of *β*_*v4*_would indicate that the drift bias towards the motivation consistent category was stronger when amygdala activation was higher. To assess if *β*_*v4*_was positive, we extracted the posterior distribution of *β*_*v4*_estimated from participants’ data and computed the proportion of the distribution that was greater than 0. This proportion denotes the probability the amygdala activation moderated motivational effects on the drift rate.

Within the same model, the starting point was also allowed to vary trial-by-trial as a function of the motivation consistent category and the amygdala activation at motivation cue onset:

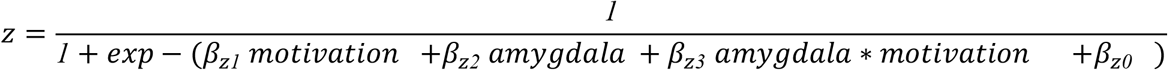

Here, the coefficient for the *amygdala*motivation* interaction reflects the extent to which the effect of motivation on the starting point depended on amygdala activation. A positive value of *β*_*z3*_would indicate that the starting point is more strongly biased towards the threshold of the motivation consistent category when amygdala activation was higher. To assess if *β*_*z3*_ was positive, we extracted the posterior distribution of *β*_*z3*_ estimated from participants’ data and computed the proportion of the distribution that was greater than 0. This proportion denotes the probability the amygdala activation moderated motivational effects on the starting point.

Model convergence was assessed using the Gelman-Rubin 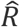 statistic. 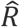 of all parameters were less than 1.004, suggesting that there were no issues with model convergence. Model convergence metrics, posterior means and 95% credible intervals of model parameters are reported in Table S3.

### Conditional Response Functions

We simulated choice and response time data from a model where the motivational bias in the starting point was dependent on amygdala activity (*amygdala bias starting point* model), and a model where the motivational bias in drift rate was dependent on amygdala activity (*amygdala bias drift* model). The simulated data reflect the pattern of choice and response time data if participants’ behavior were perfectly described by the model. Each participant was simulated performing 1000 trials at 50% scene with a model parameterized with the best fit parameters of that participant. Trial-wise amygdala activity was simulated by drawing randomly from a normal distribution with mean and standard deviation determined by the empirical mean and standard deviation of amygdala activation at the time of the motivation cue for that participant.

We plotted conditional response functions (CRF) for each model to illustrate the distinctive effect of an amygdala-dependent bias in the starting point and an amygdala-dependent bias in the drift rate. For each simulated participant, we divided trials into “fast” and “slow” trials based on a median split of RT, and “high amygdala activity” trials and “low amygdala activity” trials based on a median split of amygdala activity. We then computed the average proportion of trials on which the model predicted “more scene”, separately for face and scene motivation trials, fast and slow trials, and high amygdala activity and low amygdala activity trials. Thus, these CRFs allow us to examine the relationship between responses, motivation and RT separately for trials with high vs. low amygdala activity.

We then plotted the CRFs for the empirical data for comparison. We restricted this analysis to the subset of trials with images at 50%, as there were an insufficient number of trials at the other levels of percentage scene to be reliably divided into the 8 bins necessary for a CRF (i.e. scene vs. face motivation x fast vs. slow trials x high vs. low amygdala activity). The average proportion of trials on which participants responded “more scene” was calculated separately for each bin.

To statistically assess the effects of amygdala activity on choice and response time distributions, we ran a linear mixed effects model assessing the three-way-interaction between motivation, RT and amygdala activity on responses (LME Model M4):

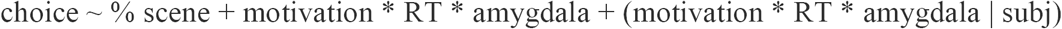

where *choice* denotes whether the participant reported that the image contained more face or more scene (face = 0, scene = 1). The LME model allowed us to treat RT and amygdala activity as continuous variables, and also allowed us to control for the effect of percentage scene on responses. As such, we were able to include trials at all levels of percentage scene for this analysis. To visualize this interaction, we fit separate LME models to estimate the effect of motivation on responses for fast vs. slow trials and high vs low amygdala activity trials.

### Confidence Analyses

We used a linear mixed effects model to examine the relationship between amygdala activity and confidence ratings, controlling for stimulus uncertainty (LME Model M5):

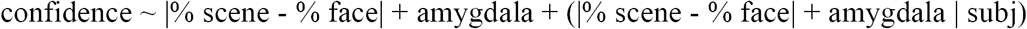

where |% scene - % face| denotes the absolute difference between % scene and % face in an image. |% scene - % face| is high when the stimulus uncertainty is low (e.g., 100 % scene and 0% face), and low when the stimulus uncertainty is high (e.g., 50% scene and 50% face). Full model specification and parameter estimates are reported in Table S1.

### Preregistration

We pre-registered how we defined the amygdala and occipitotemporal ROIs. We had also pre-registered the following hypotheses: greater amygdala activity would be associated with (1) an increased tendency to make motivation consistent categorizations, (2) stronger motivation enhancement of desirable perceptual features in the sensory cortex, and (3) stronger motivational biases in evidence accumulation in favor of the desirable category estimated using the DDM. We also pre-registered a fourth prediction that individual differences in amygdala activation would be associated with individual differences in motivational bias. However, given recent work questioning the reliability of across-subject brain-behavior correlations with a sample size of ∼30 participants (Grady et al. 2021), we no longer report this analysis. The pre-registration is available at: https://osf.io/z6xjd?view_only=030ad17d15724af2bb5f6f02ca5b34e7

## Results

Thirty participants were scanned using fMRI while they were presented with visually ambiguous images consisting of a face image and a scene image superimposed with one another. Participants had to categorize whether they saw more face or more scene in each image, and received a monetary reward for each correct categorization. At the beginning of each trial, we motivated participants to see one category over the other with a cue indicating that they would win extra money if the image was of one category (the “desirable” percept) and lose some of their earnings if the image was of the other category (the “less desirable” percept) (see *Materials and Methods*). The bonus or loss depended on the objective category of the image (i.e. whether the image actually contained more face or more scene) and not participants’ categorizations. As such, participants would earn the most money from the task if they categorized the images accurately and were not influenced by what they were motivated to see. In earlier work, we had shown that participants were nevertheless more likely to categorize the images as the desirable percept, and that this bias was associated with enhanced neural activity selective to the desirable percept in the ventral visual stream (Leong et al. 2019).

### Amygdala activity was associated with motivational bias in perceptual decisions

We divided trials into trials where participants categorized the image as the desirable percept (motivation consistent trials; e.g., categorizing an image as more scene when motivated to see more scene) and trials where they categorized the image as the less desirable percept (motivation inconsistent trials; e.g., categorizing an image as more scene when motivated to see more face). We then ran a GLM that contrasted neural activity on motivation consistent trials against that on motivation inconsistent trials at the time of the motivation cue (i.e. when participants were first informed which category would win them more money on that trial) and extracted the z-statistic in a bilateral amygdala ROI for each participant (Fig. 2A). The average *z*-statistic of the motivation consistent > motivation inconsistent contrast was significantly greater than 0 in the amygdala ROI (*t(*29*) = 2*.*45, P = 0*.*0208;* Fig. 2A, 2B), indicating that amygdala activity was higher on trials when participants made motivation consistent categorizations than when they made motivation inconsistent categorizations. Amygdala activation was not significantly associated with participants’ confidence ratings (*t*(17) = -1.61, *P* = 0.126, *b* = -0.04, SE = 0.03).

**Figure 2.**
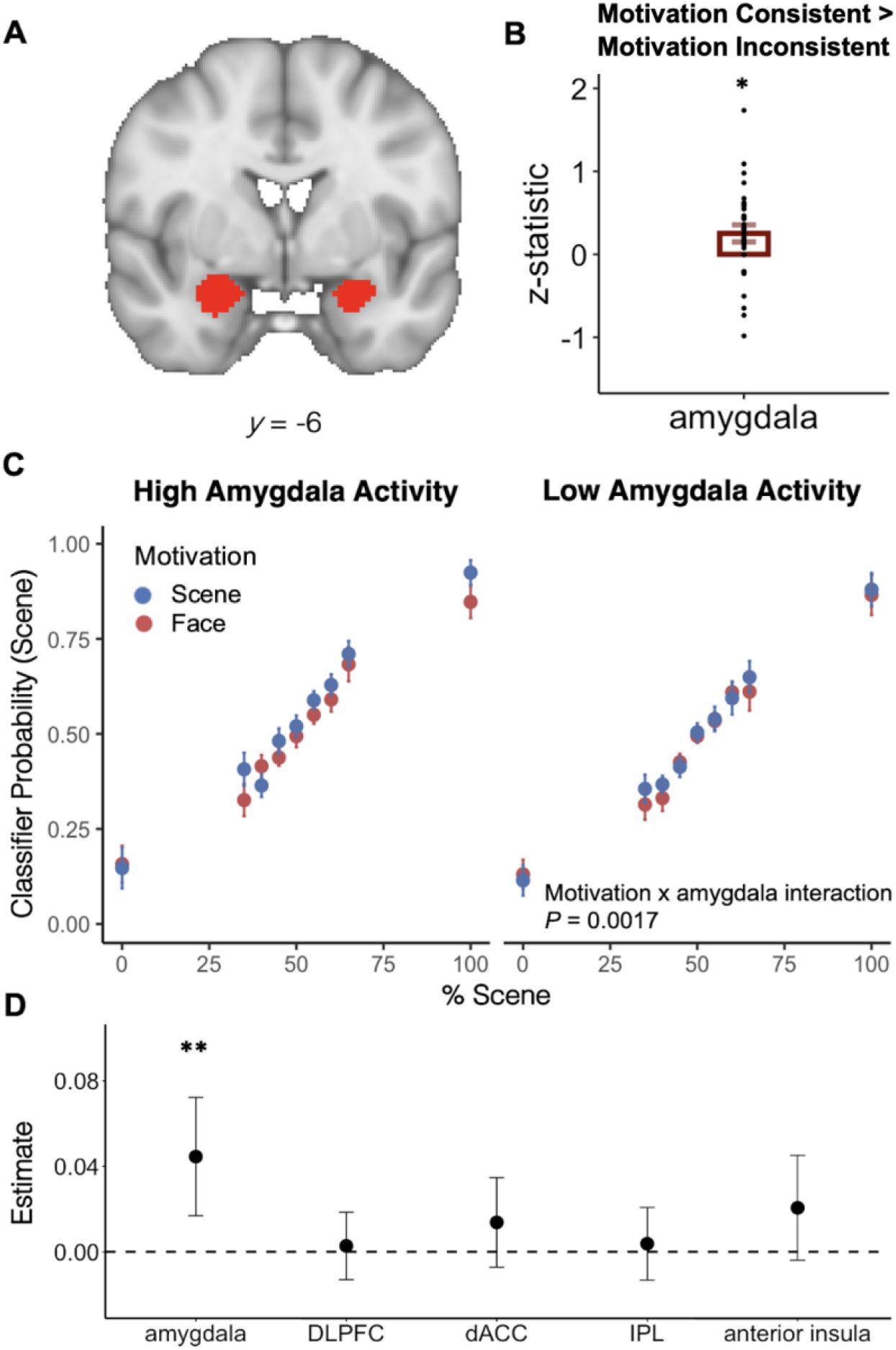
Amygdala activity was associated with motivational biases in perceptual decisions and category-selective neural activity. **A**. Amygdala ROI defined using the Harvard-Oxford Subcortical Structural Atlas. **B**. Amygdala activity was higher when participants made motivation consistent categorizations. Data points denote average z-statistic of the motivation consistent > motivation inconsistent contrast in the amygdala ROI for individual participants. **C**. Classifier probability that the participant was seeing a scene rather than a face based on the pattern of activity in the occipito-temporal ROI. On trials with high amygdala activity, category-selective activity was higher for the category participants were motivated to see. On trials with low amygdala activity, motivation had no effect on category-selective activity. **D**. Regression coefficients of the motivation x activity interaction on classifier probability for the amygdala, DLPFC, dACC, IPL and anterior insula. Error bars indicate SEM. ** p < 0.01.

### Amygdala activity was associated with stronger motivational enhancement of category-selective neural activity

The preceding analyses indicated amygdala activity was associated with motivational biases in perceptual decisions. Was amygdala activity also associated with the motivational enhancement of sensory representations? To answer this question, we used multivariate pattern analysis to quantify face- and scene-selective activity in the occipitotemporal cortex. We first trained a logistic regression classifier on BOLD data from a localizer task to categorize whether participants were seeing a face or a scene based on the pattern of activity in a occipitotemporal cortex ROI (see Methods). We then applied the trained classifier to data from the experimental task to measure the level of face and scene-selective activity on each trial.

In previous work, we had found that motivation enhances sensory representations in the occipitotemporal cortex - for a given image, face-selective activity was higher when participants were motivated to see more face, while scene-selective activity was higher when participants were motivated to see more scene (Leong et al. 2019). Here, we test if this motivational enhancement was dependent on amygdala activity. We modeled each trial as a separate regressor in a GLM.

Each regressor was created by convolving an impulse response at the time of the motivation cue with a HRF. This allowed us to estimate activation in response to the motivation cue for each trial. The resulting beta weights were then averaged within the amygdala ROI as an estimate of the amygdala activation to the motivation cue for that trial. We then used a LME model to examine the relationship between category-selective activity, motivation and amygdala activity. There was a significant interaction between amygdala activity and motivation on category-selective activity in the occipitotemporal cortex (*t*(692) = 3.15, *P* = 0.002, *b* = 0.045, *SE* = 0.014). To unpack this interaction, we divided each participant’s data into trials with high and low amygdala activity. Motivation enhanced category-selective activity on trials with high amygdala activity (*t*(30) = 2.17, *P* = 0.038, *b* = 0.032, *SE* = 0.015), but not on trials with low amygdala activity (*t*(30) = 0.43, *P* = 0.67, *b* = 0.007, *SE* = 0.016; Fig. 2C), indicating that motivational enhancement of sensory representations was dependent on amygdala activity.

Prior work implicates a network of frontoparietal regions in biasing perceptual decisions, including the dorsolateral prefrontal cortex (DLPFC), the dorsal anterior cingulate cortex (dACC), and the intraparietal lobule (IPL) (Domenech and Dreher 2010; Summerfield and Koechlin 2010; Mulder et al. 2012, 2014; White et al. 2012). In a series of exploratory analyses, we tested if these regions were associated with the motivational enhancement of sensory representations. We also considered the role of the anterior insula, which previous work had found to be involved in the accumulation of sensory evidence during perceptual decision-making (Ho et al. 2009). Activity in these regions did not moderate motivational effects on category-selective activity (DLPFC: *t*(54) = 0.34, *P* = 0.735, *b* = 0.003, *SE* = 0.01; dACC: *t*(36) = 1.29, *P* = 0.206, *b* = 0.014, *SE* = 0.011; IPL: *t*(38) = 0.44, *P* = 0.665, *b* = 0.004, *SE* = 0.010; anterior insula: *t*(37) = 1.64, *P* = 0.109, *b* = 0.021, *SE* = 0.013; Fig. 2D).

### Amygdala activity was specifically associated with motivational biases in evidence accumulation

The drift diffusion model (DDM) provides a formal framework to examine the decision processes underlying perceptual decision-making (Forstmann et al. 2016). The model assumes that perceptual decisions arise from the noisy accumulation of sensory evidence from the external environment. When the level of evidence reaches a decision threshold, the corresponding decision is made. The starting point and rate of evidence accumulation, as well as the decision thresholds and time not related to decision processes (i.e. *non-decision time*, e.g., time needed to translate the decision into a motor response), are determined by model parameters that can be estimated by fitting the model to participants’ data (Fig. 3A). In previous work, we had found that motivation biases perceptual decisions by (1) shifting the starting point towards the decision threshold of the desirable category, thereby reducing the amount of sensory evidence needed to make a motivation consistent response, and (2) by biasing sensory evidence in favor of the desirable category (Leong et al. 2019). To understand the amygdala’s contribution to motivation biases, we assessed if either or both of these biasing mechanisms were dependent on trial-by-trial fluctuations in amygdala activity.

**Figure 3.**
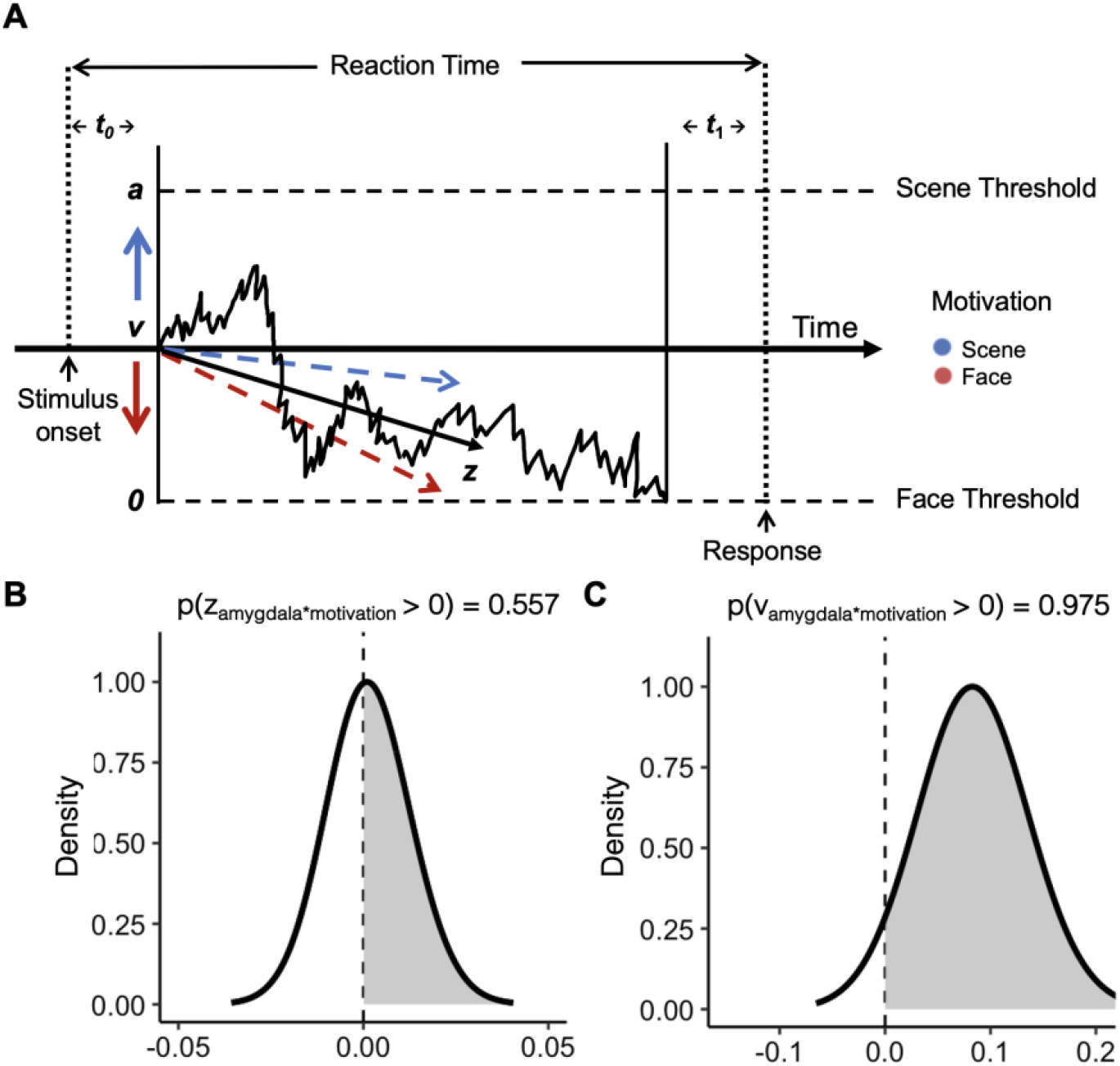
Modeling Results. **A**. Schematic diagram of the drift diffusion model (DDM). The DDM assumes that perceptual decisions arise from the stochastic accumulation of sensory evidence over time towards one of two decision thresholds. *z*: starting point, *v*: drift rate, *a*: decision threshold, *t0*: sensory delay, *t*_*1*_: time needed for response execution, the sum of *t*_*0*_ and *t*_*1*_ constitutes non-decision time. Motivation can bias perceptual decisions by biasing both the starting point and drift rate (red and blue lines). **B**. Posterior distribution of the regression coefficient of the amygdala x motivation interaction on the starting point. The distribution is centered around 0, indicating that motivational effects on the starting point did not depend on amygdala activity. **C**. Posterior distribution of the regression coefficient of the amygdala x motivation interaction on the drift rate. A large proportion of the distribution was greater than 0, indicating the motivational effects on the drift rate were dependent on amygdala activity.

We took a linear regression approach to examine the effects of the amygdala on motivational biases in the starting point and rate of evidence accumulation. Specifically, the starting point (parameter *z*), was allowed to vary as a function of the motivation consistent category, amygdala activation at the time of the cue, and the interaction between the two (amygdala x motivation interaction; see Methods). The regression coefficient on the interaction term reflects the extent to which motivational effects on the starting point depended on amygdala activity. A positive coefficient would indicate that the starting point is more strongly biased towards the decision threshold of the motivation consistent category on trials with higher amygdala activity. When we fit the model to participants’ behavioral data, the posterior distribution of the regression coefficient of the interaction term was centered around zero (*P*(*z*_*amygdala*motivation*_>0) = 0.557, mean = 0.001, 95% credible interval = -0.016 to 0.020; Fig. 3B), indicating that motivational effects on the starting point did not depend on amygdala activity.

In the same model, we allowed the rate of evidence accumulation (i.e. *“drift*” rate, parameter *v*) to vary as a function of the percentage scene in the image, the motivation consistent category, amygdala activation at the time of the cue, and the interaction between the motivation consistent category and amygdala activation at the time of the cue. Here, the regression coefficient on the interaction term reflects the extent to which motivational effects on the drift rate depended on amygdala activity. A positive coefficient would indicate a stronger bias in evidence accumulation in favor of the motivation consistent category on trials with higher amygdala activity. When we fit the model to participants’ behavioral data, the posterior distribution of the regression coefficient of the interaction term indicated a high probability that the coefficient was positive (*P*(*v*_*amygdala*motivation*_>0) = 0.975, mean = 0.084, 95% credible interval = 0.001 to 0.170; Fig. 3C). These results indicate that on trials with high amygdala activity, evidence accumulation was biased in favor of the motivation consistent category, resulting in an increased bias towards making motivation consistent responses. To visually assess model fit, we compare the distributions of model-predicted and empirical response times in Figure S2.

In exploratory analyses, we fit a model that included inter-trial variability parameters for the drift rate, starting point and non-decision-time, which allows for random trial-by-trial variability in model parameters. Consistent with our earlier results, we found that trial-by-trial amygdala activity moderated motivational biases in the drift rate (*P*(*v*_*amygdala*motivation*_> 0) = 0.991, mean = 0.14, 95% credible interval = 0.029 to 0.255) but not the starting point (*P*(*z*_*amygdala*motivation*_> 0) = 0.24, mean = -0.01, 95% credible interval = -0.034 to 0.017; Fig. S1), suggesting that our results could not be explained by random trial-by-trial fluctuations in model parameters.

### The relationship between amygdala activity and response time distributions is consistent with a motivational bias in drift rate

A bias in drift rate and a bias in starting point increases the proportion of motivation consistent responses, but have different effects on the shape of response time distributions (White and Poldrack 2014; Fig. 4). In particular, a bias in starting point has stronger effects early on in a trial than later in a trial, and would result in stronger motivational bias on fast trials than slow trials. In contrast, a bias in the drift rate has a constant effect throughout the trial, and would result in a strong motivational bias (i.e. more motivation consistent than motivation inconsistent responses) on both fast and slow trials. We can distinguish between a bias in drift rate and a bias in starting point on conditional response functions (CRF), where response proportions are plotted separately for fast (RT < median) and slow (RT > median) trials.

**Figure 4.**
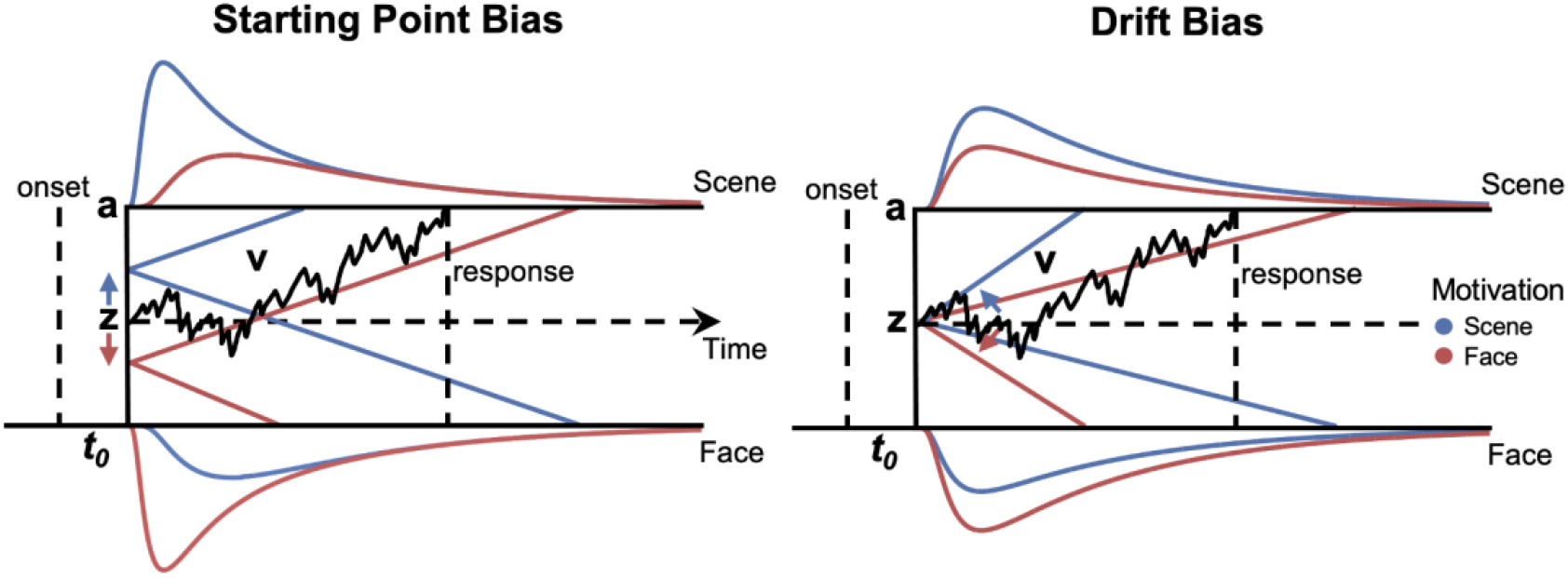
Motivational biases in the starting point and drift rate have distinguishable effects on response time distributions. The gray lines show an example trajectory of evidence accumulation on a single trial. The blue and red lines show the mean drift and resulting RT distributions when participants are motivated to see more scene and more face respectively. A bias in the starting point has a stronger effect earlier on in a trial, and would give rise to RT distributions with a stronger right-skew. In contrast, a bias in the drift rate is constant throughout a trial, and would thus scale the RT distributions proportionally such that the overall shape of the distribution does not change (right). *z, v, a, t*_*0*_ are the same as in Fig. 2. *t*_*1*_ was omitted for brevity.

To better understand the relationship between amygdala activity and model parameters, we simulated choice and response time data from models where the motivational bias in either the starting point (*amygdala bias starting point* model) or the drift rate (*amygdala bias drift* model) was dependent on amygdala activity. For each model, we plot the CRF separately for trials with high (i.e. higher than median) and low (i.e. lower than median) amygdala activity. In the *amygdala bias starting point* model, the motivational bias in drift rate, which affects both fast and slow trials, is independent of amygdala activity. Thus, motivation had a strong effect on responses on both fast and slow trials regardless of the level of amygdala activity (Fig. 5A). In contrast, for the *amygdala bias drift* model, motivation had a strong effect on both fast and slow trials when amygdala activity was high, but only a weak effect on slow trials when amygdala activity was low (Fig. 5B). This is because when amygdala activity was low, motivational effects in the model were driven predominantly by a bias in the starting point, which has stronger effects early on in a trial, and motivational effects would thus be more prominent on fast trials.

**Figure 5.**
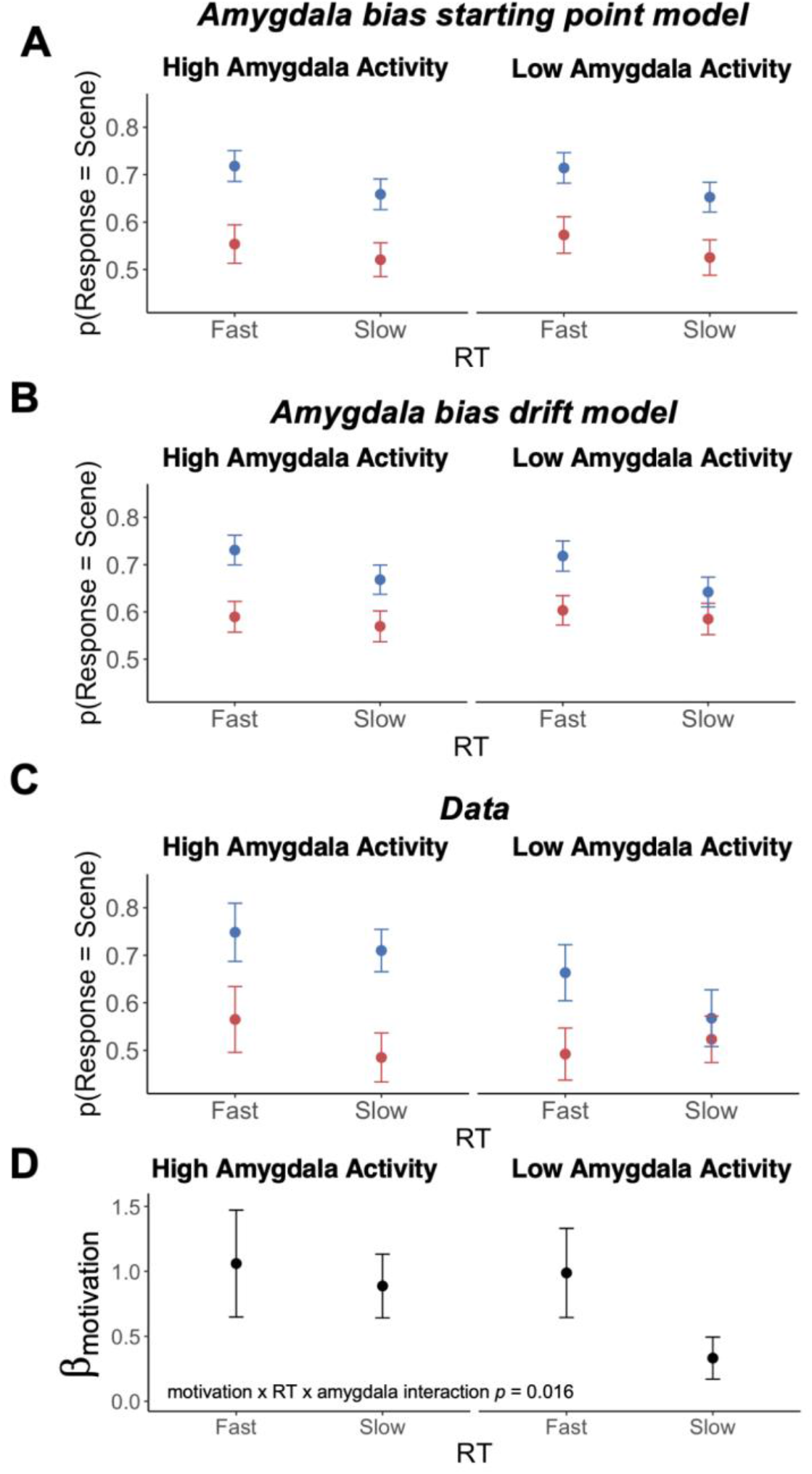
Amygdala activity moderates the relationship between motivation, RT and responses. CRF plots of data simulated from a model where amygdala biased the **A**. starting point or **B**. the drift rate. **C**. CRF plots of empirical data from trials at 50% scene. There were insufficient trials at the other levels of percentage bins to be divided into 8 bins. **D**. Regression coefficient of motivation on responses separately for fast and slow trials, and for trials with high or low amygdala activity. There was a significant three-way interaction between motivation, RT and amygdala activity such that motivational biases in perceptual decisions were weaker on slow trials with low amygdala activity.

We then assessed which pattern of response times was more consistent with the empirical data. To plot empirical CRFs, we restricted our analysis to trials at 50% scene, as that was the only level of percentage scene with a sufficient number of trials per participant to be divided into 8 bins (Fig. 5C). The empirical CRFs were most consistent with the predictions of the *amygdala bias drift* model in that when amygdala activity was high, motivation biased both fast and slow trials, but when amygdala activity was low, motivation had a weaker effect on slow trials than fast trials. Next, we sought to statistically assess the relationship between amygdala activity, response time and motivational bias in responses across the entire dataset (i.e. across all levels of percent scene). Specifically, we ran a generalized linear mixed effects model (GLMM) to test the motivation (i.e. motivated to see face vs. scene) x RT x amygdala activity interaction on participants’ responses (face vs. scene), controlling for the percentage scene of each image. Both RT and amygdala activity were treated as continuous variables in this analysis.

There was a significant three-way interaction (z = 2.41, *P* = 0.016, *b* = 0.58, SE = 0.24), indicating that the interaction between motivation and RT on responses depended on amygdala activity. To visualize this interaction, we again performed a median split based on amygdala activity to identify trials with high amygdala activity and trials with low amygdala activity. For each level of percent scene, we then performed a median split on RT to identify fast trials and slow trials. We then ran separate glmer models to assess the effect of motivation on responses separately for fast vs. slow trials, and for trials with high vs. low amygdala activity. The coefficients indicate that the three-way interaction was indeed driven by a weaker effect of motivation on responses on slow trials when amygdala activity is low, consistent with the predictions of the *amygdala bias drift model* (Fig. 5D). Together, these analyses complement the formal model-fitting results by providing a descriptive account of why the data is best explained by a model where motivational effects on the drift rate are dependent on amygdala activity.

## Discussion

Psychophysical studies have previously shown that stimulus features that have been repeatedly paired with reward tend to “capture” attention and are preferentially processed by the visual system (Anderson et al. 2011; Le Pelley et al. 2015). Relatedly, fMRI studies have found that reward value modulates sensory-related neural activity, including activity in both early visual areas (Serences 2008; Anderson 2017) and activity in the object-selective ventral temporal cortex (Hickey and Peelen, 2015; Barbaro et al., 2017; Leong, Radulescu et al., 2017). These findings highlight the pervasive influence of reward on sensory representations in the brain. Building on this body of work, we recently demonstrated that using financial incentives to motivate participants to see a particular interpretation of an ambiguous image enhances neural activity encoding stimulus features associated with said interpretation (Leong et al. 2019). This selective enhancement biases participants towards seeing perceptual outcomes that they are motivated to see. The neural mechanisms driving the selective enhancement of desirable representations, however, remain unclear.

In the current work, we show that the motivational enhancement of desirable percepts was dependent on amygdala activity. In particular, we found that heightened amygdala activity was associated with participants categorizing an image as the category they were motivated to see. Amygdala activation was also associated with stronger motivational enhancement of desirable sensory representations in the occipitotemporal cortex, suggesting that the amygdala biases perceptual decisions by driving enhanced processing of desirable perceptual features. To understand the decision computations affected by the amygdala, we fit a DDM to participants’ behavioral data and trial-by-trial amygdala activity. The model fits suggest that amygdala-dependent motivational biases on perceptual decisions were specifically related to the faster accumulation of sensory evidence in favor of desirable percepts. Together, our findings provide converging behavioral, neural and modeling evidence of the amygdala’s role in biasing perceptual decisions towards desirable perceptual outcomes.

Historically, the amygdala has been associated with the processing of fear and threat-related stimuli (LeDoux, 2000, c.f. Visser et al., 2021). More recent perspectives have emphasized a broader role of the amygdala in processing motivationally salient information, including information relevant for predicting and obtaining rewards (e.g., Pessoa and Adolphs, 2010; Pourtois et al., 2013). For example, activity in the amygdala has been shown to encode the appetitive value of stimuli (Gottfried et al. 2003; Jenison et al. 2011; Lichtenberg et al. 2017; Shanahan et al. 2021). In a series of studies with non-human primates, Peck and colleagues have also found that amygdala activity tracks spatial attention towards stimuli associated with reward (Peck et al. 2013; Peck and Salzman 2014a, 2014b). Our findings are consistent with this broader view of amygdala function that considers its role in processing reward-relevant information. Notably, our results suggest that the amygdala is not only sensitive to the value of reward-relevant stimuli, but also flexibly enhances the neural representation of what is valuable at a given moment, which in turn gives rise to a perceptual bias towards percepts that are desirable.

The amygdala can enhance sensory representations both via direct connections to sensory areas (Freese and Amaral 2005; Gschwind et al. 2012), as well as via indirect pathways involving frontoparietal regions and brainstem neuromodulatory systems (Lim et al. 2009; Pourtois et al. 2013). Recent theories have proposed that the amygdala works in concert with the locus-coeruleus norepinephrine (LC-NE) system to selectively amplify the representation of physically and affectively salient stimuli (Markovic et al. 2014; Mather et al. 2016). Building on these theories, we hypothesize that wanting to see a percept imbues affective salience to the associated perceptual features. The amygdala signals this affective salience by the selective enhancement of neural activity encoding those features. At the same time, the amygdala recruits the LC which then releases NE into the cortex. NE released in sensory areas increase the gain of sensory neurons selective to salient stimuli, such that they are more likely to fire to target inputs (Berridge and Waterhouse 2003). In addition, NE released in the amygdala further enhances the saliency signal, resulting in even greater amplification of the neural response to salient stimuli (Gallagher and Holland 1994).

Notably, amygdala activity has indeed been found to be associated with reward-driven changes in pupil dilation (Watanabe et al. 2019), a known correlate of LC-NE activity (Murphy et al. 2014; Joshi et al. 2016). Using a variant of the current task, we also recently demonstrated that motivational biases in sensory evidence accumulation were associated with increased pupil dilation (Leong et al. 2021). Thus, there is reason to hypothesize that the amygdala-dependent motivational enhancement of sensory representations observed in the current study were mediated by the LC-NE system. The small size and location of the LC present challenges to imaging the LC using fMRI (Liu et al. 2017). Recent studies have been able to take advantage of novel approaches to functionally and anatomically localize the LC (de Gee et al. 2017; Grueschow et al. 2021). Future work can use these approaches to simultaneously measure activity in the amygdala, LC and sensory regions to further characterize the neural circuitry underlying motivational biases in perceptual decisions.

Prior work examining reward-related biases in perceptual decision-making have focused on the role of frontoparietal regions (Mulder et al. 2014). Here, we did not find evidence of the involvement of frontoparietal regions in enhancing desirable sensory evidence, suggesting that our results arose from distinct biasing mechanisms. We note that in earlier studies that assign asymmetric rewards to perceptual options, participants would earn the larger reward only if they correctly categorized a stimulus as the category associated with larger reward (Summerfield and Koechlin 2010; Mulder et al. 2012). In such settings, participants would earn more reward over the course of the experiment if they biased their responses towards the category associated with larger reward (Bogacz et al. 2006; Feng et al. 2009; Fan et al. 2018). Thus, reward-related biases in these earlier experiments have largely been interpreted as a shift in response strategy to maximize reward on the task and are not necessarily related to biases in sensory evidence accumulation. Consistent with this idea, these studies have found that reward biases perceptual responses without a corresponding effect on sensory representations.

In our task, however, the additional reward associated with the motivation consistent category was not contingent on participants’ responses - participants won or lost the bonus based on the objective category of the image, regardless of their subjective categorizations. Hence, the bias observed in our task was unlikely to have resulted from a shift in response strategy to maximize reward. Instead, it was likely driven by participants wanting to see one percept over another. Using a drift diffusion model, we had previously found that this motivational bias reflects both an *a priori* response bias towards outcomes participants want to see (i.e. a bias in the starting point of evidence accumulation), and a perceptual bias in how sensory evidence accumulates over time (i.e. a bias in the drift rate) (Leong et al. 2019). The current results indicate that amygdala activity was specifically associated with the perceptual bias and not the response bias. Thus, the amygdala-dependent biases in perceptual decisions are potentially distinct from the response-related biases associated with frontoparietal regions documented in earlier studies.

In everyday life, people are rarely neutral observers indifferent to different perceptual outcomes. They are motivated to see some outcomes over others, and exhibit a bias to report seeing those outcomes over possible alternatives. The current work suggests that this bias is associated with the amygdala-dependent enhancement of desirable neural representations in sensory cortices. Together, these results expand our understanding of the neural mechanisms underlying motivational influences on perceptual decision-making. Notably, similar motivational biases have been documented across a broad array of decision-making contexts (Hughes and Zaki 2015). In one recent study, participants were asked to judge whether they were in a desirable state (i.e. an environment with greater rewards than losses) or an undesirable state (i.e. an environment with low probability of reward) by sampling evidence that was probabilistically associated with each state (Gesiarz et al. 2019). The authors found that participants accumulated evidence in a manner that was biased towards the belief that they were in a desirable state, much like the bias towards desirable visual percepts observed in our study. Could the amygdala be involved in biases in evidence accumulation beyond sensory perception? Given that the amygdala has also been implicated in value-based learning and decision-making (Jenison et al. 2011; Prévost et al. 2013; Lichtenberg et al. 2017), this is certainly possible. We believe that exploring the role of the amygdala in mediating motivational biases in other reasoning and evaluative processes would be a fruitful direction for future research.

## Acknowledgments

We would like to thank members of the Motivation and Cognition Neuroscience Lab for their feedback on this work, and Howard Nusbaum for helpful comments on an earlier version of this manuscript. Correspondence should be addressed to Yuan Chang Leong at

