## Supplementary Materials for "Trial-by-trial fluctuations in amygdala activity track motivational enhancement of desirable sensory evidence during perceptual decision-making"

### Supplementary Methods

#### Fitting a drift diffusion model with inter-trial variability parameters

We fit a drift diffusion model that also included parameters that estimated the inter-trial variability of the starting point, drift rate and non-decision time. The inclusion of these parameters allows the starting point, drift rate and non-decision time to vary randomly around a mean from trial to trial. As there usually is not enough data from each participant to meaningfully estimate these parameters on a per-subject basis, we only estimated a group-level estimate for each of these parameters. The model was otherwise identical to the model reported in the main text. Results from this model were consistent with that of the model reported in the main text, in that amygdala activity moderated motivational biases in the drift rate ($P(v_{amyg*mot} > 0) = 0.991$, mean = 0.14, 95% credible interval = 0.029 to 0.255) but not the starting point ($P(z_{amyg*mot} > 0) = 0.24$, mean = -0.01, 95% credible interval = -0.034 to 0.017; Fig. S1).

#### Posterior predictive check

To assess model fit, we generated 500 simulated datasets, each comprising the same number of participants performing the same number of trials as the real dataset. Each dataset was generated with parameter values sampled from the posterior distribution when fitting the model to participants' data. These simulated datasets reflect the pattern of choice and response time data if participants' behavior was perfectly described by the model. To compare the simulations to real data, we overlay the distribution of simulated response times with the true response time distributions, separately for face and scene responses, and separately for trials on which participants were motivated to see more face and trials on which participants were motivated to see more scene (Fig. S2). These plots serve as posterior predictive checks to assess how well the model aligns with participants' data.

**Supplementary Table 1**
**Model specification and estimated coefficients of linear mixed effects models.**

<table>
<tr><th></th><th>Formula</th><th>Term</th><th>Estimate</th><th>SE</th><th>p</th></tr>
<tr><td rowspan="5">M1</td><td rowspan="5">scene vs. face activity ~ % scene + motivation + amygdala + motivation*amygdala + (motivation + amygdala + motivation*amygdala | subj)</td><td>intercept</td><td>0.490</td><td>0.007</td><td><0.001</td></tr>
<tr><td>% scene</td><td>0.119</td><td>0.005</td><td><0.001</td></tr>
<tr><td>motivation</td><td>0.021</td><td>0.010</td><td>0.027</td></tr>
<tr><td>amygdala</td><td><0.001</td><td>0.018</td><td>0.999</td></tr>
<tr><td>motivation * amygdala</td><td>0.045</td><td>0.014</td><td>0.002</td></tr>
<tr><td rowspan="3">M2[1]</td><td rowspan="3">scene vs. face activity ~ % scene + motivation + (motivation | subj)</td><td>intercept</td><td>0.494</td><td>0.012</td><td><0.001</td></tr>
<tr><td>% scene</td><td>0.119</td><td>0.007</td><td><0.001</td></tr>
<tr><td>motivation</td><td>0.032</td><td>0.015</td><td>0.038</td></tr>
<tr><td rowspan="3">M3[2]</td><td rowspan="3">scene vs. face activity ~ % scene + motivation + (motivation | subj)</td><td>intercept</td><td>0.484</td><td>0.010</td><td><0.001</td></tr>
<tr><td>% scene</td><td>0.120</td><td>0.007</td><td><0.001</td></tr>
<tr><td>motivation</td><td>0.007</td><td>0.016</td><td>0.667</td></tr>
<tr><td rowspan="9">M4</td><td rowspan="9">choice ~ % scene + motivation * RT * amygdala + (motivation * RT * amygdala | subj)</td><td>intercept</td><td>-13.967</td><td>0.513</td><td><0.001</td></tr>
<tr><td>% scene</td><td>0.291</td><td>0.009</td><td><0.001</td></tr>
<tr><td>motivation</td><td>0.954</td><td>0.456</td><td>0.0364</td></tr>
<tr><td>RT</td><td>-0.297</td><td>0.129</td><td>0.0210</td></tr>
<tr><td>amygdala</td><td>0.479</td><td>0.334</td><td>0.152</td></tr>
<tr><td>motivation * RT</td><td>-0.109</td><td>0.181</td><td>0.547</td></tr>
<tr><td>motivation * amygdala</td><td>-0.587</td><td>0.429</td><td>0.171</td></tr>
<tr><td>RT * amygdala</td><td>-0.303</td><td>0.193</td><td>0.117</td></tr>
<tr><td>motivation * RT * amygdala</td><td>0.588</td><td>0.244</td><td>0.016</td></tr>
<tr><td rowspan="3">M5</td><td rowspan="3">confidence ~ amygdala + |% scene - % face| + amygdala + (|% scene - % face| | subj)</td><td>intercept</td><td>3.531</td><td>0.086</td><td><0.001</td></tr>
<tr><td>amygdala</td><td>-0.042</td><td>0.026</td><td>0.126</td></tr>
<tr><td>|% scene - % face|</td><td>0.018</td><td>0.001</td><td><0.001</td></tr>
</table>

Note: Models are referred to by their labels (e.g., LME Model M1, LME Model M2) in the Methods section. Formulas are written in the notation of the lme4 package in R with random effects indicated in parentheses. Variable coding - scene vs. face activity: relative strength of scene vs. face selective activity in the occipito-temporal ROI; % scene: the percentage of scene vs. face in the image; motivation: face = 0, scene = 1; amygdala: activation in the amygdala at the time of the motivational cue, choice: participants' categorizations (more face = 0, more scene = 1). [1]Data from high amygdala activity trials. [2]Data from low amygdala activity trials.

**Supplementary Table 2**
**Estimated coefficients of linear mixed effect models for additional ROIs.**

| | Formula | Term | Estimate | SE | *p* |
|---|---|---|---|---|---|
| **amygdala** | scene vs. face activity ~ motivation * amygdala + % scene + (motivation * amygdala \| subj) | intercept | 0.490 | 0.007 | <0.001 |
| | | motivation | 0.0212 | 0.010 | 0.027 |
| | | amygdala | <0.001 | 0.018 | 0.999 |
| | | % scene | 0.118 | 0.005 | <0.001 |
| | | motivation * amygdala | 0.045 | 0.014 | 0.002 |
| **DLPFC** | scene vs. face activity ~ motivation * DLPFC + % scene + (motivation * DLPFC \| subj) | intercept | 0.490 | 0.007 | <0.001 |
| | | motivation | 0.020 | 0.010 | 0.048 |
| | | DLPFC | -0.009 | 0.009 | 0.312 |
| | | % scene | 0.119 | 0.005 | <0.001 |
| | | motivation * DLPFC | 0.003 | 0.008 | 0.735 |
| **dACC** | scene vs. face activity ~ motivation * dACC + % scene + (motivation * dACC \| subj) | intercept | 0.487 | 0.008 | <0.001 |
| | | motivation | 0.028 | 0.011 | 0.034 |
| | | dACC | -0.007 | 0.011 | 0.494 |
| | | % scene | 0.120 | 0.005 | <0.001 |
| | | motivation * dACC | 0.014 | 0.011 | 0.206 |
| **IPL** | scene vs. face activity ~ motivation * IPL + % scene + (motivation * IPL \| subj) | intercept | 0.489 | 0.007 | <0.001 |
| | | motivation | 0.020 | 0.010 | 0.050 |
| | | IPL | -0.006 | 0.008 | 0.420 |
| | | % scene | 0.119 | 0.005 | <0.001 |
| | | motivation * IPL | 0.004 | 0.010 | 0.665 |
| **anterior insula** | scene vs. face activity ~ motivation *anterior insula + % scene + (motivation * anterior insula \| subj) | intercept | 0.490 | 0.008 | <0.001 |
| | | motivation | 0.026 | 0.011 | 0.022 |
| | | anterior insula | 0.008 | 0.013 | 0.542 |
| | | % scene | 0.120 | 0.005 | <0.001 |
| | | motivation * antInsula | 0.021 | 0.013 | 0.109 |

Note: Formulas are written in the notation of the lme4 package in R with random effects indicated in parentheses. Variable coding - scene vs. face activity: relative strength of scene vs. face selective activity in the occipito-temporal ROI; motivation: face = 0, scene = 1; ROI (e.g. amygdala, dACC): activity in the region of interest at the time of the motivational cue; % scene: the percentage of scene vs. face in the image.

**Supplementary Table 3**

**Priors, posterior estimates, and model convergence metrics of model parameters.**

| Parameter | Prior | Coefficient | Estimate | Gelman-Rubin $\hat{R}$ |
|---|---|---|---|---|
| *a* | $a \sim Gamma(2, 0.3)$ | $a$ | 2.264[2.166, 2.366] | 1.000 |
| *t* | $t \sim N(0.5, 0.15)$ | $t$ | 0.650[0.609, 0.694] | 1.000 |
| *z* | $z = invlogit(N(0.03, 0.04))$ | $z_{intercept}$ | 0.486[0.468, 0.505] | 1.000 |
| | | $z_{amygdala}$ | 0.007[-0.011, 0.026] | 1.001 |
| | | $z_{motivation}$ | 0.022[0.007, 0.037] | 1.000 |
| | | $z_{amygdala*motivation}$ | -0.002[-0.020, 0.018] | 1.002 |
| *v* | $v = N(0.06, 0.2)$ | $v_{intercept}$ | 0.191[0.065, 0.314] | 1.000 |
| | | $v_{stimulus}$ | 0.920[0.833, 1.010] | 1.000 |
| | | $v_{amygdala}$ | 0.087[-0.003, 0.179] | 1.000 |
| | | $v_{motivation}$ | 0.085[0.002, 0.167] | 1.000 |
| | | $v_{amygdala*motivation}$ | 0.081[0.002, 0.160] | 1.003 |

Notes: Estimates indicate the posterior means, with square brackets denoting 95% credible intervals. Model convergence was assessed using the Gelman-Rubin $\hat{R}$ statistic. $\hat{R}$ of all parameters were less than 1.004, suggesting that there were no issues with model convergence. *a*: decision threshold, *t*: non-decision time, z: starting point; *v*: drift rate, Gamma: a gamma distribution parameterized by mean and standard deviation, *N*: denotes a normal distribution parameterized by mean and standard deviation, invlogit: inverse logit transform.

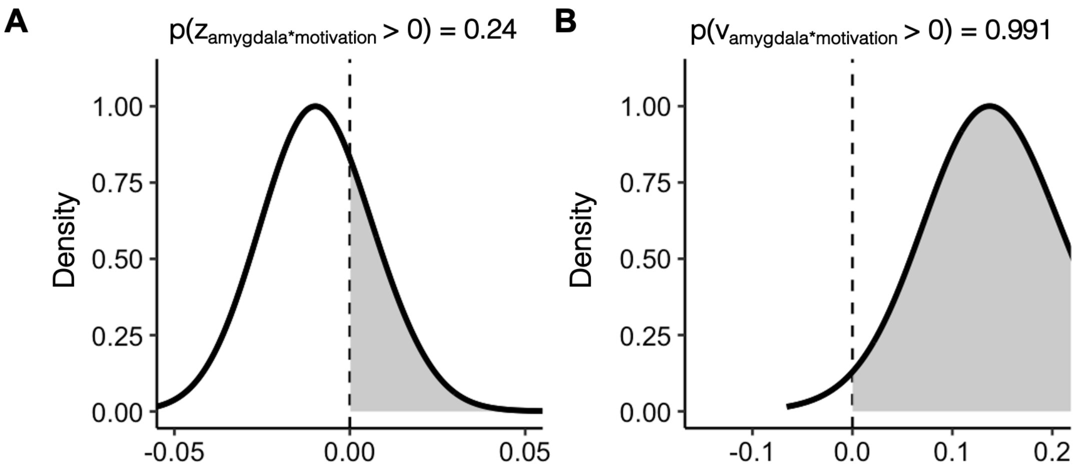


**Figure S1. Modeling Results. A.** Posterior distribution of the regression coefficient of the amygdala*motivation interaction on the starting point from a model that includes inter-trial variability parameters. The distribution is centered around 0, indicating that motivational effects on the starting point did not depend on amygdala activity. **B.** Posterior distribution of the regression coefficient of the amygdala*motivation interaction on the drift rate from the same model. The distribution skews positive, indicating the motivational effects on the drift rate were dependent on amygdala activity.

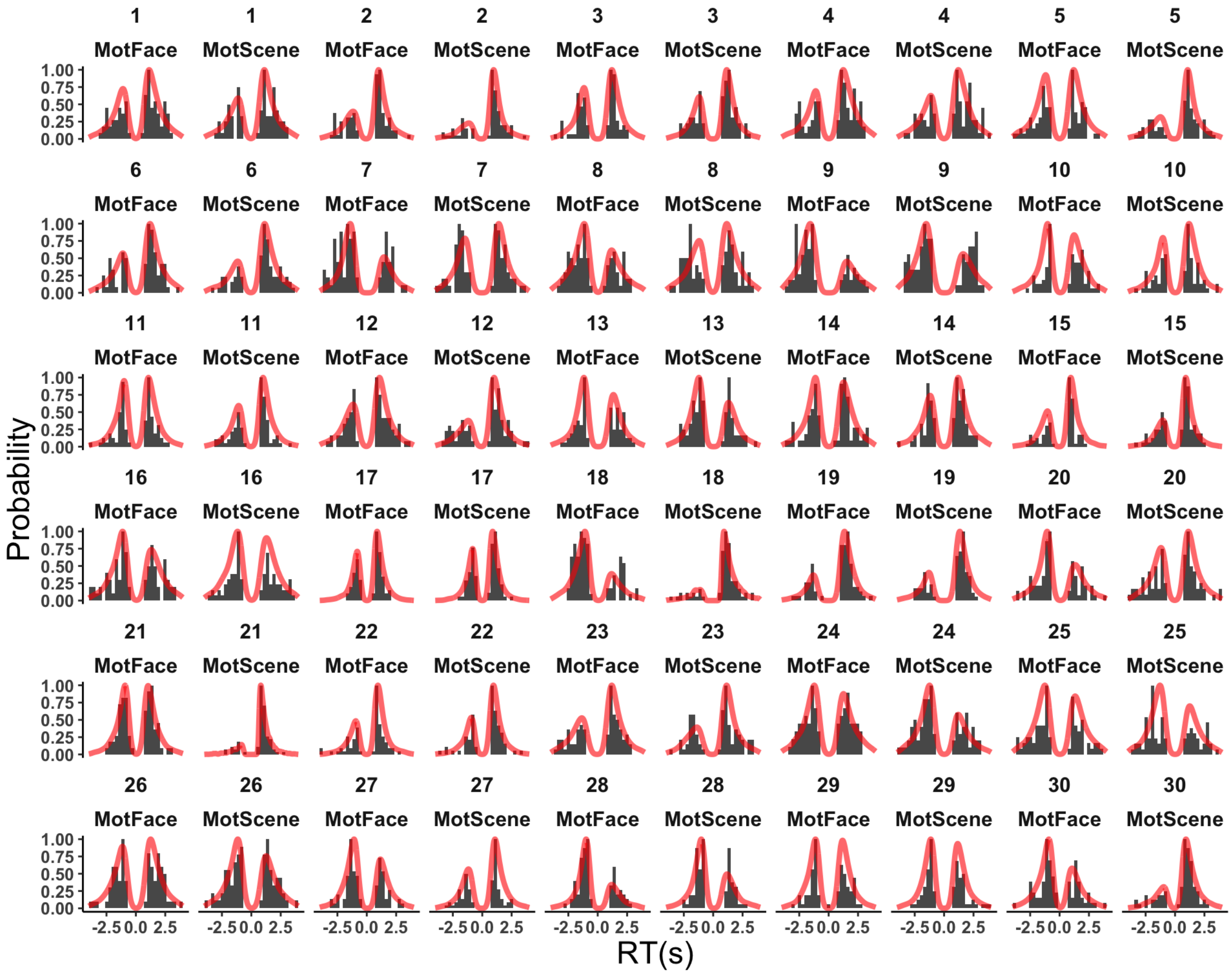


**Figure S2. Observed and predicted response time distribution of each participant (Posterior predictive plots).** The observed response time distributions for face (negative RTs) and scene responses (positive RTs) were plotted separately for each participant for when they were motivated to see more face (MotFace) and for when they were motivated to see more scene (MotScene). Red lines denote model predictions when parameterized with the best-fit model parameters. Response times for face responses were sign-flipped for illustration purposes.
